# Inferring health conditions from fMRI-graph data

**DOI:** 10.1101/295113

**Authors:** P.G.L. Porta Mana, C. Bachmann, A. Morrison

**Affiliations:** Kavli Institute for Systems Neuroscience, Norwegian University of Science and Technology, Norway; Institute of Neuroscience and Medicine (INM-6) and Institute for Advanced Simulation (IAS-6) and JARA BRAIN Institute I, Jülich Research Centre, Jülich, Germany; Simulation Laboratory Neuroscience, Inst. for Advanced Simulation, JARA, Jülich Research Centre and JARA, Jülich, Germany; Institute of Cognitive Neuroscience, Faculty of Psychology, Ruhr-University Bochum, Germany

**Keywords:** disease diagnosis, decision theory, sufficient statistics, exchangeability, parametric statistical model, schizophrenia, fMRI, Bayesian probability theory

## Abstract

Automated classification methods for disease diagnosis are currently in the limelight, especially for imaging data. Classification does not fully meet a clinician’s needs, however: in order to combine the results of multiple tests and decide on a course of treatment, a clinician needs the likelihood of a given health condition rather than binary classification yielded by such methods. We illustrate how likelihoods can be derived step by step from first principles and approximations, and how they can be assessed and selected, using fMRI data from a publicly available data set containing schizophrenic and healthy control subjects, as a working example. We start from the basic assumption of partial exchangeability, and then the notion of sufficient statistics and the “method of translation” (Edgeworth, 1898) combined with conjugate priors. This method can be used to construct a likelihood that can be used to compare different data-reduction algorithms. Despite the simplifications and possibly unrealistic assumptions used to illustrate the method, we obtain classification results comparable to previous, more realistic studies about schizophrenia, whilst yielding likelihoods that can naturally be combined with the results of other diagnostic tests.

## 1 INTRODUCTION

A 29-year-old man seeks medical advice because he finds himself in a very confused state. The clinician, after listening to the complaints of the patient, identifies some diseases that would account for the symptoms. However, the presentation is not clear cut, and treatment for some of the potential conditions have significant side effects. To come to a decision on the best course of action, the clinician decides to perform the differential diagnosis in a mathematically sound manner (Sox et al., 2013), first assigning an initial probability for the patient’s being healthy or having each of the potential conditions, taking into account age, sex, familial factors, symptoms, a psychological evaluation, the incidence of the disease, and similar prior information:

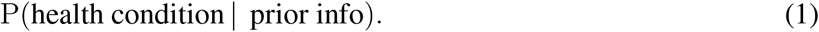

Then she orders one or more diagnostic tests to make a better informed assessment of the probabilities of the considered diseases. Among these tests she orders a structural and functional magnetic-resonance imaging (MRI) scan. The advantage of MRI lies in the non-invasive monitoring of brain structure and activity; the structural image (sMRI) is used to exclude morphological changes in the brain such as tumours, and the functional imaging (fMRI) can provide information about changes in brain activity.

With the results of the tests and of the sMRI and fMRI, the clinician updates her initial or prior probability to a “post-test” or posterior probability based on the results, according to Bayes’s theorem:

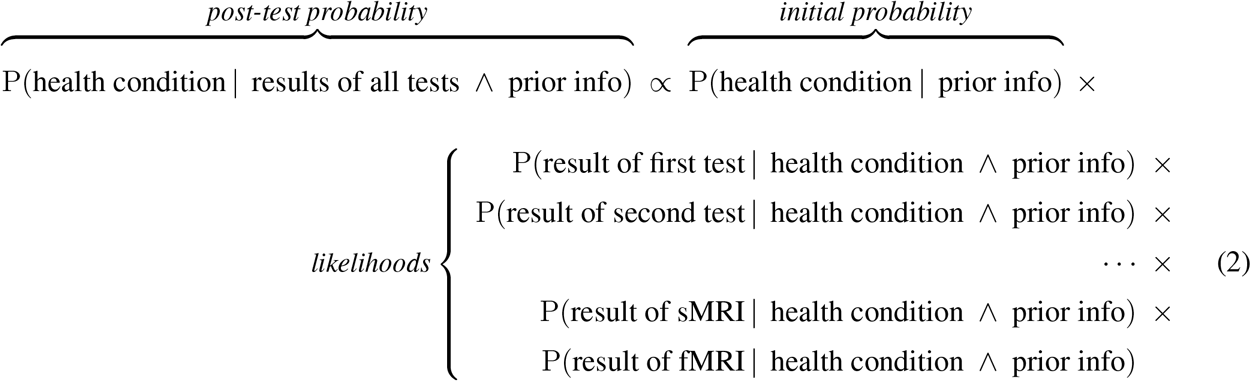

where “∧” denotes logical conjunction (“and”), and we have reasonably assumed that the result of each test does not depend on those of the other tests, i.e. that their likelihoods are independent (Jaynes, 2003, §4.2; Sox et al., 2013, §4.7).

In the update formula above, the initial probability is assessed by the clinician. To calculate the post-test probability she needs the probabilities for each test result conditional on the health condition, either “healthy” or “presumptive disease”. These probabilities are called the *likelihoods for the health condition* in view of each test. The term “likelihood” has its standard technical meaning in the present work: the probability of a proposition *A* given *B* is P(*A*|*B*), while the likelihood of *A* in view of *B* is P(*B*|*A*), i.e., A appears in the conditional (Jaynes, 2003, § 4.1; Good, 1950, § 6.1). A proposition can have high probability but low likelihood and vice versa. Probabilities, not likelihoods, are what we base our decisions upon.

The final, post-test probability is necessary to the clinician to decide upon a course of action (Sox et al., 2013, ch. 6; Goodman, 1999; Murphy, 2012, § 5.7); for example, to treat the patient according to one or another specific treatment, to dismiss him, or to order more tests. To make such decision the clinician will combine her post-test probabilities for the health conditions with a utility table (a reminder of decision theory is given in §4.3).

In the following we assume that one of the presumptive diseases the clinician has in her mind is schizophrenia. Although currently MRI does not play a role in a diagnosis of schizophrenia, there are substantial efforts to develop such analyses for this purpose (Silva and Calhoun, 2014). In this work, we focus on the diagnosis of this particular disease simply as a concrete worked example, to demonstrate how results of a diagnostic test, in this case results from function MRI imaging, can be incorporated in the diagnostic process in a principled fashion.

In short, we address the question: **how can we assign a numerical value to the likelihood**

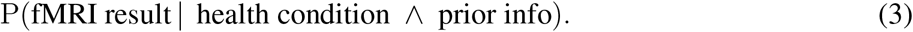

**of each health condition (‘healthy’or ‘schizophrenic’) in view of the fMRI result?** We will propose an answer that can be applied for any brain disease.

To this end it is useful to mark out some features of the approach presented so far:

I. **Modularity.** The update formula (2) combines evidence from different tests, and this combination does not need to be done at once. The clinician can multiply her initial probability by the likelihood from the first test, normalize, and thus obtain a “post-first-test” probability. Later she can multiply this probability by the likelihood from the second test, normalize, and thus obtain a post-second-test probability; and so on with any number of other tests, a number that the clinician needs not fix in advance. She can therefore store the value of the likelihood from the fMRI result, to later combine it with new likelihoods from future tests to form a new, better-informed post-test probability.
II. **Decision-theoretic character.** The clinician’s final goal is not simply a healthy/schizophrenic classification, but a *decision* upon a course of action about the patient (Sox et al., 2013, chs 6, 7; Jaynes, 2003, chs 13, 14; Raiffa and Schlaifer, 2000). This distinction is important: for example, a treatment without contraindications might be recommended even if there is only a 10% probability that the disease is present; or a dangerous treatment might be recommended only if there is a 90% probability that the disease is present. The modularity of the present approach extends to the decision stage, because the post-test probability can be used with different decisions and utilities, which can also be updated later on. For example, after beginning a treatment the clinician happens to read about a new kind of treatment, having new benefits and contraindications. Using the post-test probabilities she already has, she may re-evaluate her decision using an updated utility table that includes the new treatment.
III. **Incomplete knowledge.** In general, we lack a complete biological understanding of the relation between brain activity and the health condition under study. In this case, the likelihoods can only be assessed by relying on examples of known *health condition–fMRI data* pairs, usually called a training or calibrating dataset. Moreover, this training dataset is often very small.
IV. **High dimensionality.** The fMRI data are positive-valued vectors with 10^7^−10^8^ components or more (Lindquist, 2008). This high dimension impacts the calculation of likelihoods and probabilities.

The first two points above are great advantages of the present approach, and also the reasons why it cannot be based on machine learning algorithms for deterministic classification; such methods give the clinician a dichotomous, “healthy/schizophrenic” answer, with no associated uncertainty. This answer cannot be used by the clinician to weigh the benefits and risks of different courses of action, the assessment of which needs the probabilities of the health conditions. Probabilistic algorithms, on the other hand, are not flexible for combining evidence: they give a probability for the health condition, not a likelihood; and only the latter can be combined with the likelihoods from other tests, or stored for later reuse and combination.

We therefore approach the question of assigning the likelihoods (3) by means of the probability calculus, the same calculus from which eq. (2) is derived. What we will do is in essence no different from current Bayesian statistical analyses and modelling; but we would like to emphasize some aspects of this modelling that are usually left in the background. The probability calculus can be regarded as the extension of formal logic (truth calculus) to plausible inference (Jeffreys, 2003; Jaynes, 2003; Hailperin, 1996), a view also supported in medicine (Greenland, 1998; Maclure, 1998; Goodman, 1999), which has been proven with increasing rigour by Koopman (1940b; 1940a; 1941), Cox (1946; 1961; 1979), Pólya (1949; 1968), and many others (Horvitz et al., 1986; Paris, 2006; Halpern, 1999; Snow, 1998; Dupré and Tipler, 2009; Terenin and Draper, 2017). The derivation of a probability proceeds much like an “axioms → logic rules → theorem” derivation in formal logic: one starts from the probabilities of some propositions, and by applying the probability rules, arrives at the probability of the desired proposition, eq. (3) in our case.

We will show this procedure step by step in the case of our problem, in order to expose where assumptions and approximations enter the derivation. These may be improved by other researchers, or replaced by different ones when the method is applied to a different problem. Our discussion is inspired by Mosteller & Wallace’s (Mosteller and Wallace, 1963) brilliant, thoughtful analysis of a statistically similar problem in a very different context.

The approach we follow deals naturally with the four points listed above. The small size of the training dataset, point III. above, is not an issue because the probability calculus allows for training datasets of any size. In fact, the calculus allows us to continuously update our inferences given new training data, making our inferences more and more precise and less likely to be affected by outliers.

The unmanageable size of our data space, point IV. above, will force us to make auxiliary assumptions that will translate into the choice of a reduced data space, discussed in §2.3, and into the use of parametric statistical models, discussed in §2.4. Regarding the latter, we will emphasize that assumptions about relevant and irrelevant information in our data may translate into mathematical statistical models. It is often difficult to relate biophysical considerations about quantities measured in the brain to the shape of a probability distribution, especially in multidimensional quantities. The notion of *sufficient statistics* (Dawid, 2013; Bernardo and Smith, 2000, ch. 4; Lindley, 2008, § 5.5; Diaconis and Freedman, 1981; Cifarelli and Regazzini, 1982; Lauritzen, 1988; Kallenberg, 2005), discussed in §2.4.2, is a helpful bridge between biophysical considerations and probability distributions. The idea is that it may be easier for us to conceive a connection between biophysical considerations and some special statistics of our measurements, than between biophysical considerations and an abstract multidimensional distribution function. This “translation” is powerful, because if one finds the assumptions about relevance or irrelevance of some data unreasonable, one can then make different assumptions, resulting in a different statistical model.

Models inspired by sufficient statistics – especially their comparison and selection – can nevertheless be computationally demanding owing to the multidimensional integrals in their formulae, even when these are addressed by modern numerical methods such as Monte Carlo (MacKay, 2003, ch. IV; Murphy, 2012, chs 23–24). In the present study we shall use analytically tractable statistical models, but availing ourselves of Edgeworth’s “Method of Translation”(1898; Johnson, 1949; Mead, 1965): the simple but potentially very fertile idea of transforming a quantity into a normally distributed one, discussed in §2.4.3.

The possible combined choices of reduction of the data space, of sufficient statistics, and of transformations into normal variable, lead to a variety of possible models and likelihoods to be used by the clinician. Which is the “best” one? We discuss several criteria for choice in §2.5, settling on one based on expected utility. We also briefly discuss the remarkable observation that common Bayesian criteria based on weight of evidence and Bayes factors (Jeffreys, 2003, chs V, VI, A; Good, 1950; MacKay, 1992a; Kass and Raftery, 1995) for the fMRI data gives results opposite to those of the expected-utility criterion.

In this article, we will calculate the likelihoods for the health conditions, eq. (3), and assess the models for so doing, in the following steps. First, in §2.1, we briefly discuss schizophrenia and the use of fMRI to diagnose it, introducing a concrete dataset of fMRI data for schizophrenic and healthy patients. We then show that a simple and natural assumption, called *exchangeability*, would lead to a unique value of the likelihoods (3) if the training dataset were large enough (§2.2). However, with a small training dataset we must face two problems: unmanageably large dimensions of the data space, and the need to specify prior beliefs, also involving functions on infinite-dimensional spaces.

To solve the first problem, in §2.3 we assume that information adequate for our health inference can be found in a reduced data space of the fMRI, which we construct from time correlations between groups of voxels. To solve the second problem we introduce parametric statistical models in §2.4 using the notions of *sufficient statistics* and of transformation into normal variables, mentioned above. We discuss how these models learn from the data and select three models as possible candidates. We then consider several criteria to select one of the three models against our data, as an example, and discuss how a more realistic assessment could be made in a real application (§2.5). We conclude with a discussion (§3) on how the choice of sufficient statistics and prior probabilities could be improved, and on the relation to machine-learning methods.

Our statistical terminology and notation follow ISO standards (ISO, 2009, 2006).

## 2 RESULTS

### 2.1 Selection of clinical use case and fMRI-data acquisition

Schizophrenia is a psychiatric disorder that comprises various symptoms that are categorized into positive (e.g. hallucinations), negative (e.g. loss of motivation) and cognitive (e.g. memory impairment) disease patterns. A common disease cause for all these widespread symptoms is still unknown. Functional magnetic resonance imaging (fMRI) has been used to gain insight into modifications in functional connectivity in this disease. In the resting state, functional connectivity is measured either by asking the subject to fulfil a certain task or at rest, instructing the subject to think about nothing specific but not fall asleep. In this condition, both increased and decreased functional connectivity have been reported in the default mode network, although the hyperactivity seems to be reported more often (Hu et al., 2017). Moreover, widespread connectivity changes in the dorsal attention network and the executive control network have been detected (Woodward et al., 2011; Yu et al., 2016).

Beyond these individual sub-networks, many studies have found profound changes in macroscopic brain structures, e.g. a shrinkage of whole brain and ventricular volume, reduced gray matter in frontal, temporal cortex and thalamus, and changes in white matter volume in frontal and temporal cortex (Shenton et al., 2010; Ellison-Wright and Bullmore, 2009, 2010). Since both gray matter loss and white matter changes are found, it is reasonable to conclude that not only the intrinsic activity of single areas is modified, but also the interplay of different brain areas, in particular in frontal and temporal cortex. It has been argued that these alterations in long range connectivity are responsible for a range of disease symptoms that are not attributable to single areas (Friston and Frith, 1995). Taking this disconnect hypothesis as a starting point, we can reach the working hypothesis that these changes are also reflected in the functional activity of the brain, and that fMRI images can be used to distinguish schizophrenic from healthy patients. We therefore conclude that schizophrenia is an appropriate condition to demonstrate our approach.

We requested data of schizophrenic and healthy patients from Schizconnect^1^, a virtual database for public schizophrenia neuroimaging data. In our request we asked for resting state T2*-weighted functional (rfMRI) and T1-weighted structural magnet resonance images (MRI) from patients participating in the COBRE study either with no known disorder or diagnosed as schizophrenic according to the Diagnostic and Statistical Manual of Mental Disorders (DSM) IV, excluding schizoaffective disorders. In the COBRE study, the voluntary and informed participation of the subjects was ensured by the institutional guidelines at the University of New Mexico Human Research Protections Office. The ensuing dataset comprised 91 healthy patients and 74 schizophrenic patients. Out of these we randomly selected 54 healthy and 49 schizophrenic subjects, to permit demonstration our method on a small dataset with unequal group size. A detailed description on the exact experimental design and the MRI scanning is provided by Çetin et al.(2014).

### 2.2 Calculation of probabilities: exchangeability

Let us describe our context more precisely and set up some mathematical notation. We have:

- A number of possible health conditions, in our example healthy (H) and schizophrenic (S). The variable c denotes health condition.
- A space of possible fMRI data. They are vectors with 10^7^–10^8^ or more positive components (Lindquist,2008). The variable ***f*** denotes an fMRI result.
- A set of *n* patients, labelled in some way, the variable *i* denoting their labels. These labels may reflect information about the times the patients were examined, or about their geographical location. This possibility is important in the considerations to follow. In our study *n* = 104.
- Knowledge of the health condition and of the fMRI result of each patient. Let us use the propositions

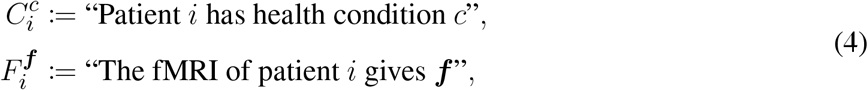

the latter to be understood within a very small interval (***f***, ***f*** + d***f***). In our study we have *n*_H_ = 55 healthy and *n*_S_ = 49 schizophrenic patients.

For brevity we denote by *C^c^* the conjunction of the propositions 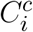 for all patients having health condition *c*, i.e. our knowledge about which patients have that health condition; and analogously for *F^c^*. By *D^c^* we denote all data about patients with health condition *c*; by *D* we denote all our data.

- An imaginary patient, labelled “0”, whose fMRI result ***f*** is known, but whose health condition is not.
- Other pre-test information, denoted by *I*; for example the clinician’s initial diagnosis of the health condition of patient 0, and the results of any other diagnostic tests.
- The probabilities 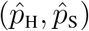 for the health condition of patient 0, conditional on the pre-test information, including the results from other tests. We call these *pre-test probabilities*. Note that they may differ from the *initial* probabilities of eqs (1)–(2), because they may include the likelihoods from other tests.
- A set of decisions about the patient 0 and their utilities conditional on the patient’s health condition. We shall simply consider two decisions: dismiss (D) or treat (T). See §4.3 for a summary of decision theory.

Our goal is to assign numerical values to the likelihoods for the health conditions (3): the conditional probability distribution that the fMRI result of patient “0” is ***f*** given the health condition of that patient and all other data. In our notation,

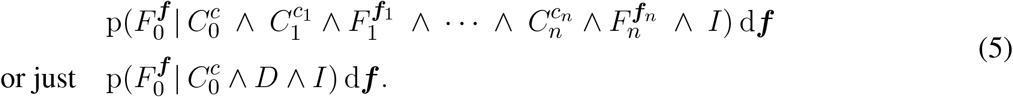

A natural assumption helps us restrict the values the distribution above may have. Within a group of patients *having the same health condition* we assume that the probability that a patient shows a particular fMRI ***f**_i_* does not depend on the particular value of the patient’s label *i*, no matter how many patients we have or may later add in that health group. This assumption is called *partial exchangeability* (de Finetti,1938;Diaconis and Freedman, 1981; Aldous, 1985; Diaconis, 1988). If the labels carry e.g. temporal or geographical information, partial exchangeability means that we do not expect to observe particular kinds of fMRI results more often in the future than in the past, or more frequently in one location than another. As a concrete example: fix three possible fMRI results ***f***_1_, ***f***_2_, ***f***_3_ (each is a vector with 10^7^–10^8^ positive components) and consider the fMRI tests of three schizophrenic patients: say, one from five years ago in Germany, one from last week in Scotland, and one to be done six months from now in Italy. Partial exchangeability means that the probability that the German patient’s test gave ***f***_1_, the Scottish’s gave ***f***_2_, and the Italian’s will give ****f***_3_*, is numerically equal to the probability that the German’s gave ***f***_2_, the Scottish’s *f*_3_, and the Italian’s will give ***f***_1_; and likewise for all six possible permutations of the three results. Keeping the same fixed fMRI results ***f***_1_, ***f***_2_, ****f***_3_*, we now consider three healthy patients instead, who may also live in different times and places. Partial exchangeability means that also in this case the values of the six possible joint probabilities obtained by permutation must all be equal – but this value can be different from the one for the schizophrenic patients considered before. Hence the term “partial”: we can freely exchange the joint results within the schizophrenic group and within the healthy group without altering their probabilities, but not across groups. This assumption extends in an analogous way to more patients.

The assumption of partial exchangeability might not be completely true when we consider geographical or epochal differences, but we may still consider it as a good approximation. We are not making any exchangeability assumptions about the probabilities of the health conditions of our patients, though, because the incidence of a disease does often change with time and can depend heavily on geographical location.

To express partial exchangeability mathematically, suppose that the patients i = 1, 2, 3, … have health condition *c* = H and the patients *i′* = 1′, 2′, 3′, … health condition *c* = S. Then the joint distribution for their fMRI results satisfies

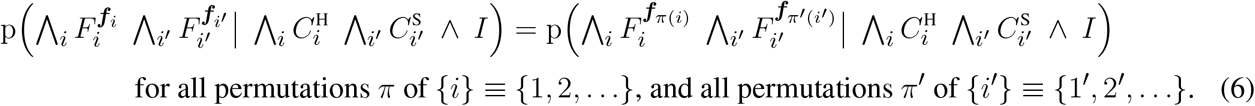

The assumption of partial exchangeability is simple and quite natural – and very powerful: it implies, as shown by de Finetti (1938; Diaconis, 1988, § 3; Bernardo and Smith, 2000, § 4.6), that the joint distributions above must have the form

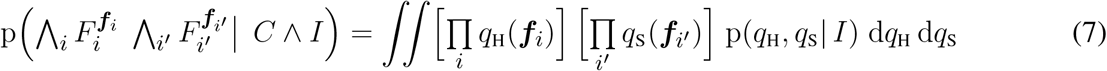

where *q*_H_, *q*_S_ are distributions over the possible values of ***f***, and p(*q*_H_, *q*_S_|*I*) is a “hyperdistribution” over such distributions, determined by the assumptions I. The double integral (which can be understood as a generalized Riemann integral:Lamoreaux and Armstrong, 1998; Swartz, 2001; Kurtz and Swartz, 2004), is over all distributions *q*_H_, *q*_S_. In other words, de Finetti’s theorem say that the joint probability distribution for the fMRIs of healthy and schizophrenic patients can be seen as the product of independent distributions, identical for healthy cases and identical for schizophrenic cases but different for the two cases, mixed over all possible such pairs of distributions with weight p(*q*_H_, *q*_S_|*I*).

As a very cursory example, suppose we want the joint probability that a healthy patient has fMRI result ***f*** and a schizophrenic one ***f′***. De Finetti’s formula first tells us to consider all possible distributions over positive vectors. As usual with infinities, “all” must be made precise by specifying a topology (for details see e.g. de Finetti, 1938; Diaconis and Freedman, 1981; Aldous, 1985; Diaconis, 1988); but intuitively these distributions comprise, e.g., multivariate truncated normals, gammas, exponentials, truncated Cauchys… and innumerable distributions that we can imagine and don’t have a specific name for; all with their possible parameter values. De Finetti’s formula tells us to choose one distribution *q*_H_, from all those possible ones, for the healthy case and one *q*_S_ for the schizophrenic case, and to attach a weight to this pair, p(*q*_H_, *q*_S_|*I*); then to calculate this pair at the values ***f***, ***f′*** and multiply them: *q*_H_ (***f***) × *q*_S_ (***f′***). Then we consider a new pair of distributions, attach a weight to them, and again multiply their values at ***f*** and ***f′***. And so on, until all possible pairs are considered. Finally we calculate the sum of all such products, weighted accordingly: 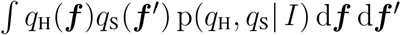.

The generalization of the formulae above to more than two health conditions, or when only one health condition is considered, is straightforward.

As the cursory example above made quite clear, an integral over probability distributions is a mathematically complicated object (cf.Ferguson, 1974) and may not seem a great advancement in assigning a value to the distribution (5). In defence of de Finetti’s formula we must say that it is completely manageable with discrete data spaces and provides a great insight in the way we reason about probability in relation to repeated events (de Finetti, 1937; Lindley and Phillips, 1976; Kingman, 1978;Koch and Spizzichino,1982; Dawid, 2013; Bernardo and Smith,2000, § 4.2). It also has several important consequences for our inference, which we now discuss.

Using de Finetti’s formula and the definition of conditional probability we can rewrite our goal plausibility (5) as

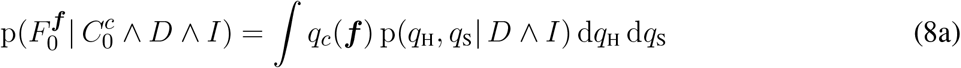

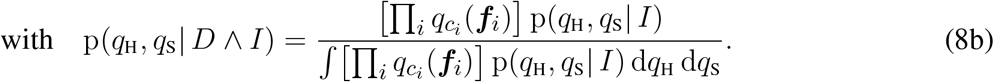

The latter is called posterior distribution since it is conditional on all data *D*.

Excluding pathological prior distributions (Diaconis and Freedman, 1986), this posterior distribution becomes more and more concentrated on two particular distributions 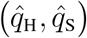, fully determined by the data, as our data *D* comprise a larger and larger number of patients. This concentration occurs independently of the original shape of the distribution p(*q*_H_, *q*_S_|*I*). In this limit our probability distribution (5) becomes

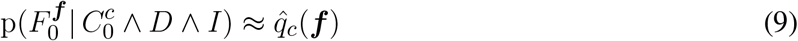

with 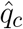 completely determined by the data *D*. This would solve our plausibility assessment (5).

De Finetti’s formula therefore tells us also the theoretical limit by which the pre-test probability for the health condition of the patient, 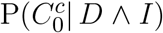, can be improved by the fMRI result. For example, if 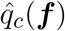 is more or less uniform in ***f*** or has the same peaks in ***f*** for each *c*, then the fMRI is of no use for discriminating the health condition of the patient. This result follows mathematically from the assumption of exchangeability (6) and the rules of probability calculus, hence no amount of ingenuity could overcome this limit.

In our case the amount of data *D* is not enough to allow the use of the approximation (9). We should use the general formula (8), but it is unwieldy in two respects. First, the fMRI result ***f*** of a patient is a positive-valued vector with 10^7^–10^8^ components or more (Lindquist, 2008), so the distributions *q*_H_, *q*_S_ are highly multidimensional. Second, the formula asks us to consider in principle *all* such distributions, as explained in the example above.

We tame this double unwieldiness in two ways. First, it is conceivable that not all information contained in the fMRI result ***f*** of a patient be relevant to discriminate the patient’s health condition *c*. The integral in eq. (8), if it could be performed, would automatically winnow out the relevant information (Jaynes, 2003, ch. 17), possibly reducing the problem to a lower-dimensional set in the space of fMRI data. Being unable to perform the integral, we must try to apply heuristics based on our understanding of the target conditions and perform such dimensional reduction by hand. For example, by employing the hypothesis that in schizophrenic patients the time correlation between brain regions is altered with respect to healthy ones. We can thus address the first problem by reducing fMRI data ***f*** to a manageable set of graph properties ***f***, and applying our inference directly on these, as explained in the next section.

Second, we entertain a working assumption about which features of our graph data {***f**_i_*} from a set of patients are relevant for inferences about new patients. We can, for example, assume that only the first and second moments are relevant for making predictions about the graph quantities to be observed in the new patients; these moments are then called *sufficient statistics*. Assumptions of this kind reduce the infinite-dimensional space all possible distributions (*q*_H_, *q*_S_) to a finite-dimensional space of a parametric family of distributions of exponential type, as explained in §2.4.

### 2.3 Trimming the data space: functional connectivity

The preprocessed functional image in standard space, in which the activity for each voxel is recorded, consists of approximately 10^17^ time series of 140 time points. We reduce this huge data space by the following steps.

First, we consider only the activity of the voxels belonging to the 94 regions defined by the lateral cortical Oxford atlas (see §2.1 and Desikan et al., 2006) and average the activity of all voxels in a region (details in §4.1). Second, we measure the functional connectivity defined by the Pearson correlation coefficient between pairs of regions, obtaining 94 × 93/2 = 4371 connectivity weights. This is still a considerable data space for our computational resources, so we select *d* = 40 connectivity weights that exhibit the greatest difference in their connection weight average across the schizophrenic and healthy groups.

The resulting distributions for four of these connectivity weights are depicted in fig. 1. Note that for each considered brain connection, the histograms of the healthy group and the schizophrenic group display significant overlap, such that none of them could be used in isolation to reliably discriminate between two groups.

**Figure 1.**
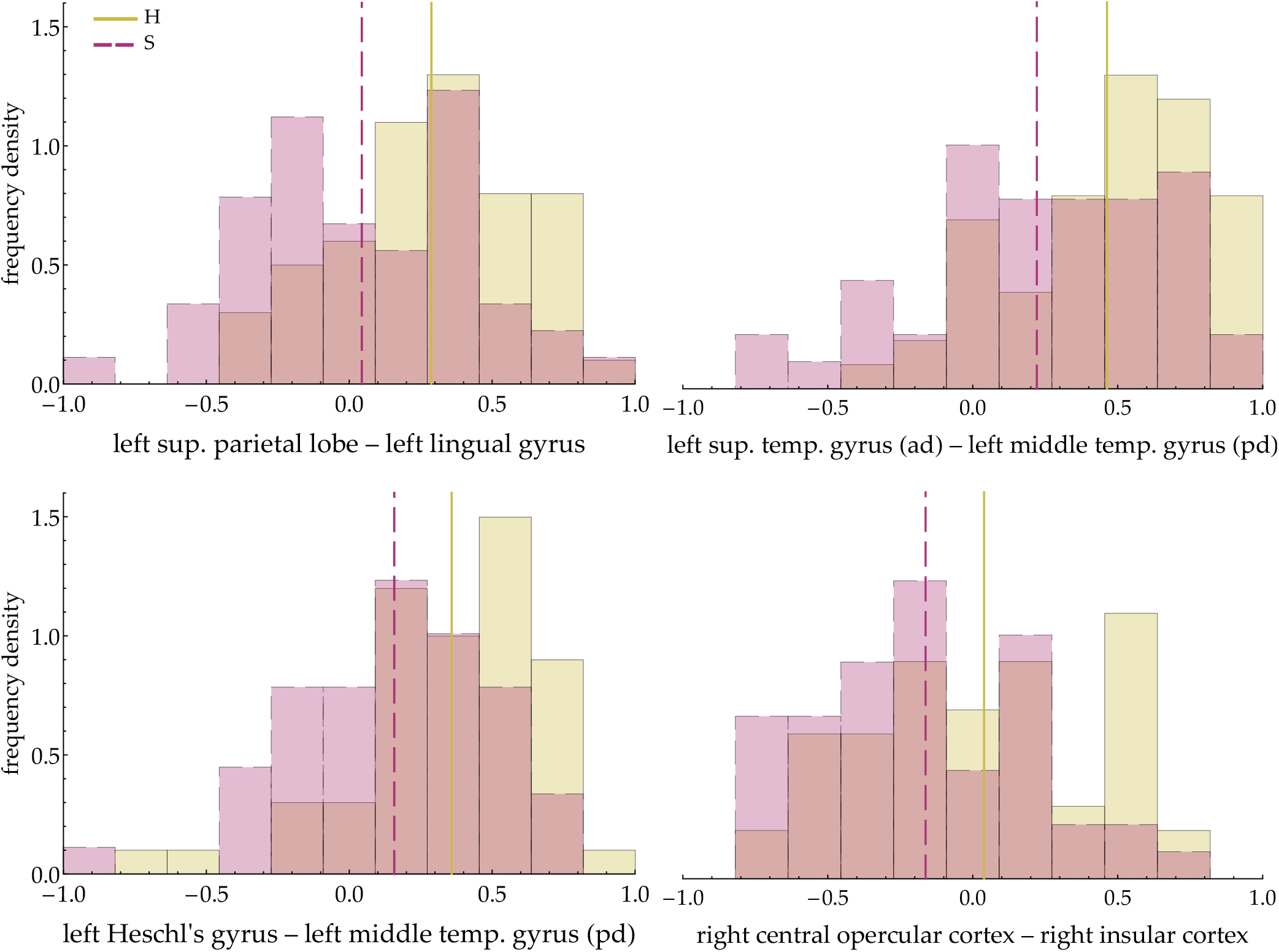
Distributions of functional connectivities of schizophrenic and healthy patients. Normalized density histograms of connectivity weights for healthy (H, yellow, solid) and schizophrenic (S, red, dashed) patients of four cortical connections selected to demonstrate our statistical framework. Empirical means are shown as vertical lines. All connectivities for healthy and schizophrenic patients have substantial overlap, evidenced by the darker regions in the histograms.

Our data space is therefore vastly reduced, from [0, ∞[^10^17^^ to [−1, 1]^40^. We let the symbol ***f*** stand for the set of connectivity weights extracted from the fMRI data, rather than the full fMRI data themselves. With this new meaning of the symbol ***f***, the likelihoods for the health conditions (5) and de Finetti’s formulae (7)–(8) remain formally unchanged, but now involve a data space with much fewer dimensions.

### 2.4 Trimming the distribution space: models by sufficiency and generalized normals

#### 2.4.1 Parametric models

The integrals in de Finetti’s formulae (7)–(8) still represent a mixture of all imaginable pairs of probability distributions (*q*_H_, *q*_S_) over the space [−1, 1]*^d^*, and are therefore extremely complex. We now examine two ways to reduce this integral to a manageable set of distributions and to obtain analytically tractable formulae.

In the Bayesian literature, the complication of considering all possible distributions **q*_c_*(***f***) of the quantity ***f*** is typically sidestepped by restricting them to a finite-dimensional set of distributions *L_c_*(***f|θ****_c_*), identified or indexed by a finite number of parameters ***θ**_c_*. For this reason, such a set is called a parametric family of distributions. An example of parametric family is the set of *d*-variate normal distributions parameterized by their mean ***μ*** and covariance matrix ***Σ***. With this restriction, the integrals in de Finetti’s formulae (7) and (8) represent mixtures of distributions within the parametric family, the weight for each distribution being represented by a weight for its parameters. That is, we are no longer considering mixtures of all possible multivariate truncated normals, gammas, exponentials, etc., as in the example of §2.2, but only mixtures of truncated normals, say, with different means and covariance matrices. These integrals are thus ordinary finite-dimensional integrals. What happens to formulae (7)–(8) is that

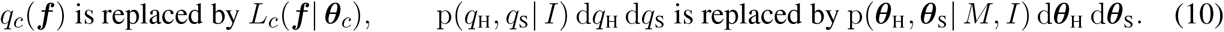

The distribution *L_c_* is called the likelihood of the parameters ***θ**_c_*, and p(***θ***_H_, ***θ***_S_ | *M*, *I*) is the prior parameter distribution. A parametric family and a prior distribution over its parameters are jointly called a parametric statistical model, which we denote by *M*. The term “model” is justly criticized by some probability theorists (see e.g. Besag & Kalman in Besag et al., 2002) but widely used, so we shall adopt it here.

In our present problem, the parametric statistical model needs not be the same for all health conditions: for example, we could use normal distributions for one condition and beta distributions for another, if that choice better reflected the distributions of connectivity weights under the two different health conditions. For this reason, we use the subscript “*c*” in the formulae above. The likelihood we want to determine, eq. (5), thus becomes, from eq. (8) with the replacements (10),

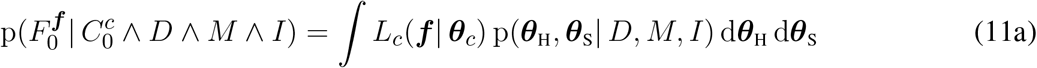

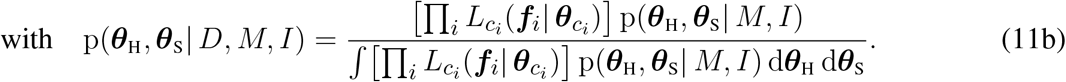

#### 2.4.2 Models by sufficient statistics

The choice of a statistical model often appears as an art and a matter of experience. Notable statisticians have voiced concerns over the esoteric character of this choice. Dawid (1982a, p. 220) says: “Where do probability models come from? To judge by the resounding silence over this question on the part of most statisticians, it seems highly embarrassing”. And Diaconis (1988, § 8, p. 121) remarks:

> de Finetti’s alarm at statisticians introducing reams of unobservable parameters has been repeatedly justified in the modern curve fitting exercises of today’s big models. These seem to lose all contact with scientific reality focusing attention on details of large programs and fitting instead of observation and understanding of basic mechanism. It is to be hoped that a fresh implementation of de Finetti’s program based on observables will lead us out of this mess.

Authors like these have also tried to develop intuitive methods to choose a model, for example by proving that a parametric family can be uniquely determined by some symmetry assumptions about our inferences, or by other information-theoretical properties (Bernardo and Smith, 2000, ch. 4; Lindley, 2008, § 5.5; an enlightening discussion of this topic is given by Dawid, 2013). In the present study we want to emphasize, as Cifarelli & Regazzini (1982) did, that a statistical model can be chosen by selecting a *sufficient statistics* (Kolmogorov, 1942; Freedman, 1962; Diaconis and Freedman, 1980, 1981; Cifarelli and Regazzini, 1982; Lauritzen, 1988; Diaconis, 1992; Kallenberg, 2005, and the textbook references above). Here is an example.

Imagine that we have patients labelled *i′* ∈ {1′, 2′, …}, and n patients labelled *i* ∈ {1, 2, …}, all with the same health condition. Of the second set of patients we also know the connectivity weights {***f**_i_*} obtained by fMRI. We want to specify the joint probability distribution p({***f**_i′_*}|{***f**_i_*}, *I*) that the fMRIs of the patients {*i′*} yield connectivity weights {***f**_i′_*}, conditional on our knowledge of the connectivity weights of the *n* patients {*i*}. Now assume that the probabilities for the fMRI results are exchangeable, so that de Finetti’s formulae (7) and (8) hold. Also assume that in order to specify the joint distribution we do not need the full set of data {***f**_i_*}, but only their number n and some statistics, e.g. the empirical mean and covariance matrix of these data,

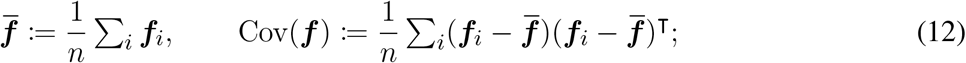

the rest of the details of the data {***f***_i_} being irrelevant. In other words, we are assuming that the statistics above are *sufficient* for us to make predictions as if we had the full data. In symbols,

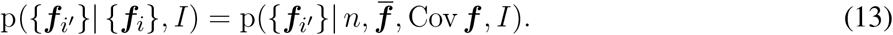

If this is true no matter the number of patients {*i′*} and {*i*}, then these statistics are called (*predictive) sufficient statistics*. There are several notions of sufficiency, including the traditional one by Fisher (1922) and Neyman (1935), but they all are more or less equivalent (Bernardo and Smith, 2000, § 4.5.2).

The assumption of the existence of sufficient statistics has a very important consequence for de Finetti’s formulae (7) and (8): the space of possible prior distributions is hugely reduced, constrained to be non-zero only over a parametric family of distributions that is determined by the sufficient statistics. The replacement (10) takes place, leading to the simpler formula (11) for the likelihoods of the health conditions. The number of parameters is equal to that of the sufficient statistics. The proof of this reduction was given by Pitman and Koopman (Koopman, 1936; Pitman, 1936; Darmois, 1935; for generalizations, e.g. to the discrete case, see Hipp, 1974; Andersen, 1970; Denny, 1967; Fraser, 1963; Barankin and Maitra, 1963). When the sufficient statistics are the mean and covariance matrix, as above, the likelihoods turn out to be (truncated) multivariate normal distributions.

In the slightly more complicated case of two or more health conditions and the assumption of partial exchangeability (6), this theorem leads to formula (11) with likelihoods *L_c_* determined by the sufficient statistics that we have chosen for the different health conditions (Bernardo and Smith, 2000, § 4.6). The prior distribution over the parameters of the likelihoods, p(***θ***_H_, ***θ***_S_ 11), is not determined by the theorem, but has to be determined by other consideration that can again involve symmetry and information theory.

As we mentioned in the introduction, the notion of sufficient statistics can be a helpful bridge between biophysical considerations and the specification of probabilities. It may be easier to conceive and understand a connection between biophysical considerations and some statistics of our measurements, than between biophysical considerations and an abstract multidimensional distribution function. Once such statistics are selected, they in turn uniquely select a probability distribution for us. Vice versa, if a statistical model based on some sufficient statistics proves to be very reliable in its predictions, we may conclude that its sufficient statistics must have an important biological meaning.

In the rest of this study we shall use three statistical models determined by three different sufficient statistics. Our choice of statistics is unfortunately not biologically motivated, as such models have yet to be determined for fMRI data. However, they are adequate to demonstrate the approach and we hope that authors with more experience will pursue this line of thought and derive better-motivated sufficient statistics.

#### 2.4.3 Edgeworth’s “method of translation”: generalized normal models

The assumption of a sufficient statistics makes the integrals in de Finetti’s formulae (7) and (8) finite-dimensional, but these integrals and other expressions that depend on them, like the post-test probabilities, may still lack a closed form and be analytically intractable. In this case we could use numerical methods, a computationally costly possibility we consider in the Discussion, §3.3. In the present work we choose models with closed-form formulae instead; their swift calculation facilitates the model comparison to be illustrated later.

Our starting point is an analytically tractable model by sufficient statistics that has been the subject of much study (Gelman et al., 2014, § 3.6; Minka, 2001; Murphy, 2007): it has a normal likelihood, with mean **λ** and covariance matrix ***Λ*** as parameters, and a normal-inverse-Wishart prior distribution over these parameters. This parameter prior maintains the same functional form when updated with training data; this kind of prior is called *conjugate*(DeGroot, 2004, ch. 9; Diaconis and Ylvisaker, 1979). This model is outlined more in detail in §4.2.

We try to make the normal + normal-inverse-Wishart model more flexible by combining it with an idea that Edgeworth (1898) called “Method of Translation”, discussed also by Johnson (1949) and Mead (1965): the transformation of a quantity into a normally distributed one. That is, instead of considering the connectivity weights ***f***, we consider transformed quantities *l*(***f***), where l is a component-wise monotonic function, and suppose the latter quantities to be normally distributed. This leads to generalized-normal likelihoods of the form

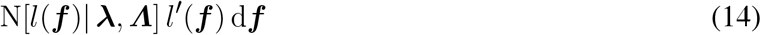

where N is the normal distribution with mean **λ** and covariance matrix ***Λ***, and *l′* is the Jacobian determinant of *l*.

This simple idea has an amazing scope: it allows us to explore a vast range of non-normal likelihoods – in particular likelihoods defined on bounded domains such as [−1, 1]*^d^* – and to keep the low computational costs of the conjugate prior. In our present problem it has also an additional convenient feature: in the calculation of the post-test probability, the Jacobian determinant *l′* disappears, as eq. (19) below shows.

These generalized-normal models, which we denote by *M_l_*, are also determined by a choice of sufficient statistics, analogous to eq. (12): the mean and covariance matrix of the transformed data {*l*(***f**_i_*)},

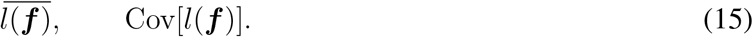

In our partially exchangeable case, with formulae (11), we need to specify a likelihood *L_c_* for each health condition *c*. We assume these two likelihoods *L*_H_, *L*_S_ to be functionally identical generalized normals, i.e. the function *l* is the same for the healthy and the schizophrenic case; but their means and covariance matrices (**λ**_H_, **Λ**_H_) and (**λ**_S_, **Λ**_S_) can be different.

We also need to specify a joint parameter prior for (**λ**_H_, ***Λ***_H_; **λ**_S_, ***Λ***_S_). To use the analytic advantage of the conjugate prior, we assume that the distribution for these parameters is a product of independent distributions:

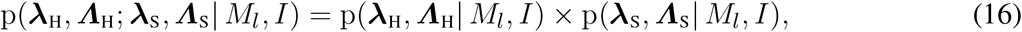

each of them being a normal-inverse-Wishart distribution described in §4.2. This independence assumption is quite strong and has an important consequence: *the likelihood for a health condition only depends on the data from previous patients having that same health condition*.

With the assumptions above, the likelihood for the health condition needed by the clinician has a closed form for this model (see §4.2). The likelihood for patient 0’s being healthy, in view of the patient’s measured connectivity weights f, and given the data (***f**_i_*) from previous *n*_H_ healthy patients, is

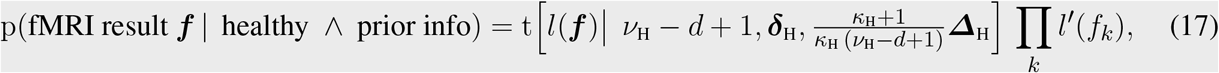

where t is a multivariate t distribution with *ν*_H_ − *d* + 1 degrees of freedom, mean ***δ***_H_, and scale matrix 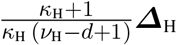. This distribution has covariance matrix 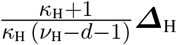 (note the different denominator from the scale matrix), and approaches a generalized normal for large *ν*_H_. The final factor is the Jacobian determinant of *l*.

The most important feature of this likelihood is the dependence of the coefficients (*κ*_H_, ***δ***_H_, *ν*_H_, ***Δ***_H_) on the data (***f**_i_*) of the previous *n*_H_ healthy patients:

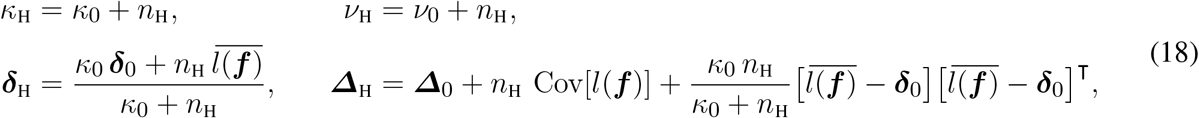

where (*κ*_0_, ***δ***_0_, *ν*_0_, ***Δ***_0_) are prior coefficients that represent the clinician’s knowledge before any patients were observed. As the number *n*_H_ of observed healthy patients increases, the probability for the transformed data *l*(***f***) tends to a normal distribution with mean and covariance matrix equal to the empirical average and covariance matrix of the transformed data. The formulae above show that previous data enter only through the sufficient statistics 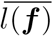 and Cov[*l*(***f***)].

An analogous formula holds for the likelihood for the patient’s being schizophrenic, with coefficients (*κ*_S_, ***δ***_S_, *ν*_S_, ***Δ***_S_) that depend on the data of previous schizophrenic patients and some initial coefficients. The function *l* and the prior coefficients (*κ*_0_, ***δ***_0_, *ν*_0_, ***Δ***_0_) could be different for the healthy and schizophrenic cases, but for simplicity we assume them identical for both health conditions.

If 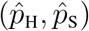 is the pre-test probability distribution for the health condition of patient 0, his post-test probability to be healthy is

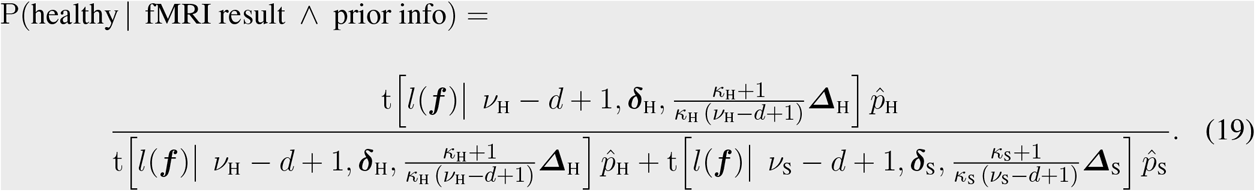

Note that the Jacobian determinants *l′* do not appear in this formula.

#### 2.4.4 Generalized normal models in our study

In the rest of our study we compare three different transformations l of the connectivity weights f, with one set of prior coefficients (*κ*_0_, ***δ***_0_, *ν*_0_, ***Δ***_0_) each:

**Logit-normal model:** A slightly modified logit transformation

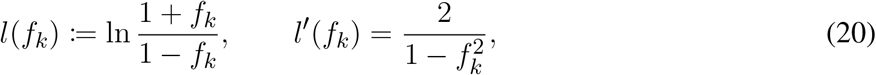

with prior coefficients

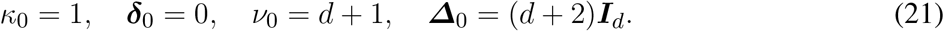

**Tangent-normal model:** A tangent transformation

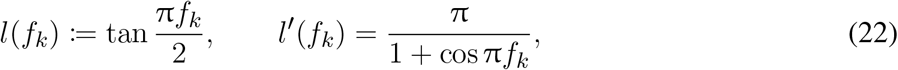

with prior coefficients

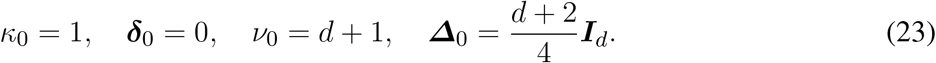

**Normal model:** An identity transformation (that is, no transformation at all)

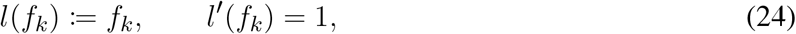

with prior coefficients

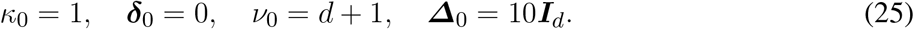

For brevity we shall denote *l*(***f***):= (*l*(*f_i_*)).

The first two transformations, plotted in the upper panel of fig. 2, map the bounded domain] − 1, 1[ of the connectivity weights into the reals, and thus restrict the generalized-normal likelihood to meaningful values of the connectivity weights. The last model instead allows for connectivity weights outside their meaningful bounds. It can be conceived as the model of a person who has no precise knowledge of what the quantities ***f*** are. Since the clinician’s final predictions concern health conditions given data ***f***, not the data f themselves, this model can still be meaningfully used. The probabilities for the connectivity weight *f_i_* conditional on the prior coefficients above are shown in the lower panel of fig. 2.

**Figure 2.**
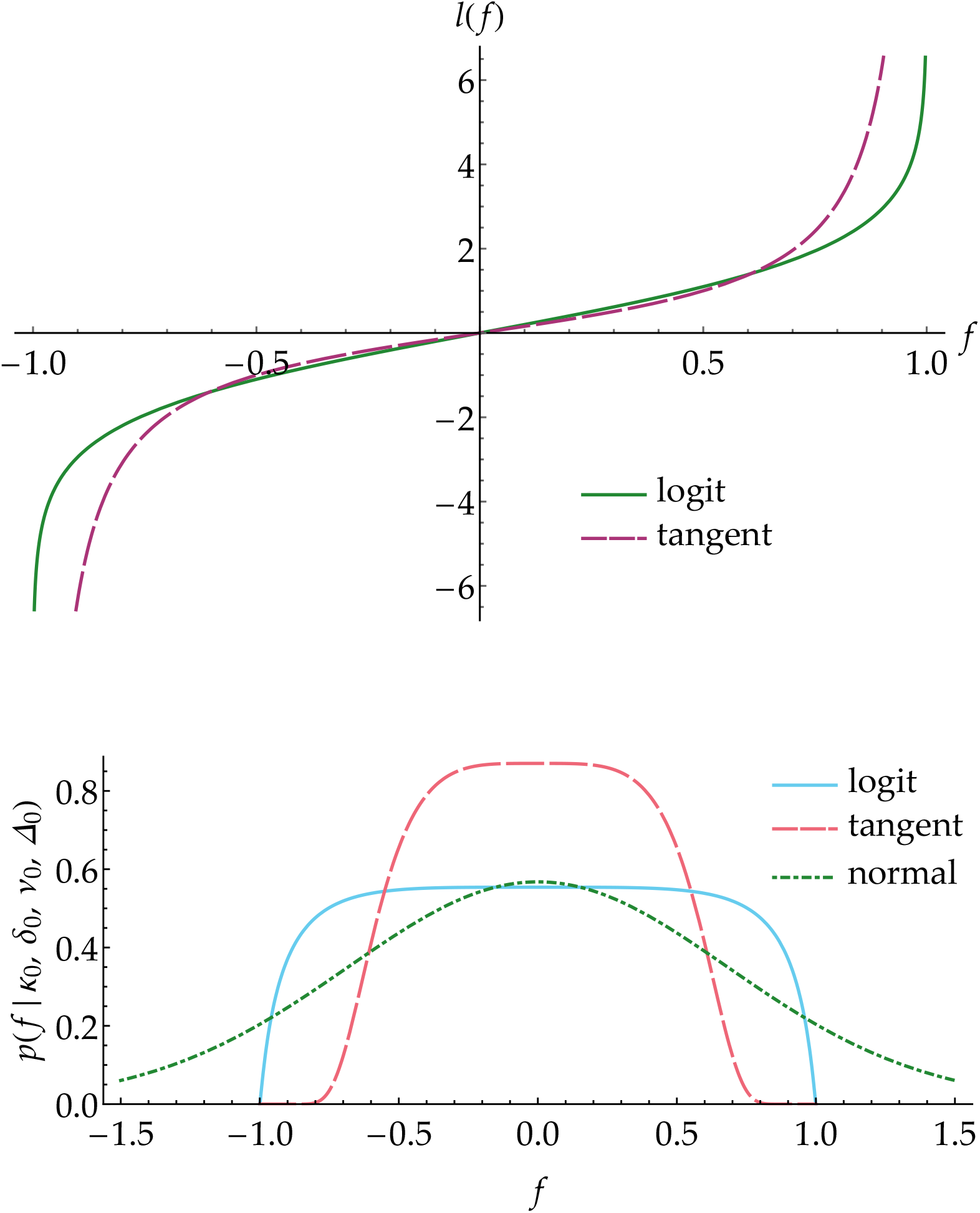
General normalized models. Upper panel: Generalized logit and tangent transformation functions as defined in the text. Lower panel: Prior distributions for any connectivity weight f conditional on the three generalized normal models defined in the text

The prior coefficients are chosen by the following criteria: uniform marginal distributions of the correlations between connectivity weights (leading to *ν*_0_ = *d* + 1 as explained before); large uncertainty in the location parameters (*κ*_0_ = 1); symmetry with respect to the origin (***δ***_0_ = 0); a prior distribution for the connectivity as flat as possible (its second derivative vanishes at the origin, leading to the values of ***Δ***_0_ above). In the case of the identity transformation we have chosen a ***Δ***_0_ that somewhat concentrates the prior around the true range of the connectivity weights, [−1, 1].

The numerical values of the main quantities used throughout this study are summarized in table 1.

**Table 1.** Numerical values in our study

|  |  |
| --- | --- |
| $n_H = 55$ | healthy patients |
| $n_S = 49$ | schizophrenic patients |
| $d = 40$ | graph parameters |
| $\kappa_0, \delta_0, \nu_0, \Delta_0$ | prior coefficients: see eqs (21), (23), (25) |

### 2.5 Model comparison and selection

#### 2.5.1 Criteria for model comparison

Any two statistical models differ in two main characteristics: their predictive power and their learning speed. Predictive power is a model’s capacity to give high probability to propositions that turn out to be true, during and especially after its learning phase (cf.Dawid, 1982b). Learning speed is how quickly a model reaches unchanging, stable predictive probabilities as it gets updated with new data; note, however, that a model may also never reach stable probabilities (see e.g.Bruno, 1964; Berk, 1966); “Alas, this seems like a model of the way things work in practical inference – as more data comes in, one admits a richer and richer variety of explanatory hypothesis”(Diaconis, 1988, § 3, p. 113). Thus, our choice of a model depends on the relative importance we give to these two characteristics.

These two characteristics need not go hand in hand: a model can quickly learn with very little data but settle on probabilities with poor predictive power; conversely, it can reach great predictive power but only after a long learning phase with a huge amount of data. A model can also “unlearn”, i.e. its predictive power initially increases and then decreases before stabilizing. The latter phenomenon can happen because every model initially makes its prediction through a mixture of likelihoods – the integral (11), in our case the t distribution (36) – but as the training continues the mixture is replaced by a single likelihood, in our case a generalized normal (14), as explained in §2.4.3. It may happen that the mixture of likelihoods has higher predictive power than a single likelihood, and in this case a decline in predictive power will be observed. A model with such behaviour is obviously unfit for the clinician’s goal; this also means that the sufficient statistics on which it is based capture very poorly the differences in connectivity weights between health conditions.

When only a small amount of training data is available, it can be difficult to assess which of two models has or will have the greater predictive power. The first model can initially reach a greater predictive power than the second, but the second model may eventually reach greater predictive power than the first, with further training data. Models having the same likelihood, however, have the same final predictive power; their learning speeds depend on their parameter priors.

In a diagnostic problem like that faced by our clinician, the choice of a model is ultimately dictated by the predictive power of the post-test probabilities given by the model; but if we have little training data, the learning speed of the model is also of some importance. Several quantitative criteria can be conceived to assess these two characteristics:

1. the post-test probabilities the model gives to the correct health conditions for all training data, i.e. P(*c*_1_, *c*_2_, …);
2. the post-test probabilities the model gives to the correct health conditions in the final phase of the training only, i.e. P(*c*_last_|*c*_1_, *c*_2_, …);
3. the expected utility the model yields for all training data;
4. the expected utility the model yields in the final phase of the training only.

Post-test probabilities (criteria 1, 2) are important for obvious reasons, and utilities (criteria 3, 4) are important because the clinician’s overall problem is one of decision, as emphasized in the Introduction. Consideration of all data (criteria 1, 3) is important if we are interested in the performance of the model for the whole set of patients; but consideration of the final data only, conditional on the previous ones (criteria 2, 4), tells us how much the model has learned (compare with a similar remark in model comparison using Bayes factors by Berger and Pericchi, 1996).

We must keep in mind that these criteria assess a statistical model not by itself but in combination with other factors, since they also depend on pre-test probabilities (which can be influenced by other diagnostic tests) or utilities.

Applied to our models, each of these four criteria gives a very similar picture. We shall calculate the results for criteria 2 and 4, the latter with two different utility tables. This calculation can be explained in very intuitive terms:

Imagine that the *n*_H_ = 55 healthy and *n*_S_ = 49 schizophrenic patients visit the clinician in turn, in an unknown order. For each patient, let us further assume that the clinician has pre-test probabilities 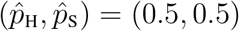, i.e. she is completely uncertain about the patient’s health condition. The incidence in the population is much lower than 50%, of course, but the patients presenting themselves for diagnosis are not representative of the full population.

As stated in §2.2, pre-test probabilities represent the clinician’s uncertainty before the fMRI test is made; they can be based for example on a first diagnosis considering symptoms and medical history of the patient and of the patient’s family, on psychological evaluations, and on other diagnostic tests. Here we assume complete uncertainty for demonstration purposes.

The clinician acquires the fMRI result f for that patient, and uses the statistical model, trained with the data from all the patients that previously visited her, to update to a post-test probability for schizophrenia *p*_S_, given by eq. (19); obviously *p*_H_ = 1 − *p*_S_. Now the clinician must make a decision – say, treat or dismiss – based on the expected utilities of the decisions available. Each decision has a different utility depending on the patient’s true health condition, as summarized by a table. We consider two tables: a symmetric one

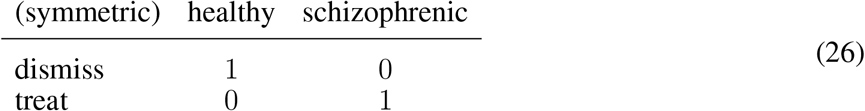

and an asymmetric one

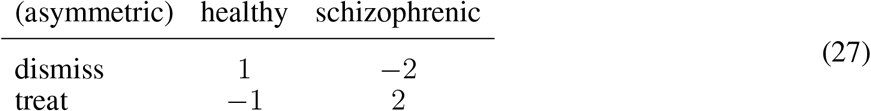

In order to maximize the expected utility, as explained in §4.3, the clinician dismisses the patient if *p*_S_ < 1/2 in the case of the symmetric utility table, and if *p*_S_ < 1/3 in the case of the asymmetric one; and treats the patient otherwise. After the clinician’s decision is made, the patient’s true health condition is revealed and we record the actual utility gained for each table, and the post-test probability the clinician assigned to the true health condition, or its logarithm, usually called “negative surprise”(Bartlett, 1952; Good, 1956, 1957a). The health condition and fMRI data of this patient are used to update the model. The next patient is received, and so on, until all patients have been examined.

The particular sequence of utilities and log-probabilities recorded in the manner just described depends on the exact order by which the 104 patients visit the clinician. We take an approximate average over all possible 104! ≈ 10^166^ orders by randomly sampling 520 000 of them. The values of these averages for the final patient constitute the quantitative criteria 2 and 4.

With the symmetric utility table (26), the average utility is also the average number of schizophrenic patients for which the model yields *p*_S_ > 1/2. The average utility for the last patient, when the model has been trained with the rest of the patients, is therefore a form of leave-one-out cross-validation (Allen, 1974; Stone, 1974; Kotz et al., 2006, vol. 2, pp. 1454–1458). The asymmetric table (27), slightly more realistic, tells us that dismissing a schizophrenic patient has worse consequences than treating a healthy one, and treating a schizophrenic patient has better consequences than dismissing a healthy one (McKenzie, 2014; Ho et al., 2000). For this reason the patient is dismissed, more conservatively, only if *p*_S_ < 1/3. Note that scaling a utility table by a positive factor or shifting its values by a constant represent changes in the unit of measure and in the zero of utilities, and therefore do not affect our relative comparison of the statistical models.

#### 2.5.2 Results for our three models

The averaged sequences of utilities and log-probabilities calculated as in §2.5.1 are shown in fig. 3. The R code for the calculation is publicly available (Porta Mana et al., 2018). The results, summarized in table 2, are qualitatively identical by the two criteria and two utility tables we chose: the normal model gives the best values for the final patient, followed by the logit-normal model; the tangent-normal model has the worst final predictive power, at or below chance level.

**Figure 3.**
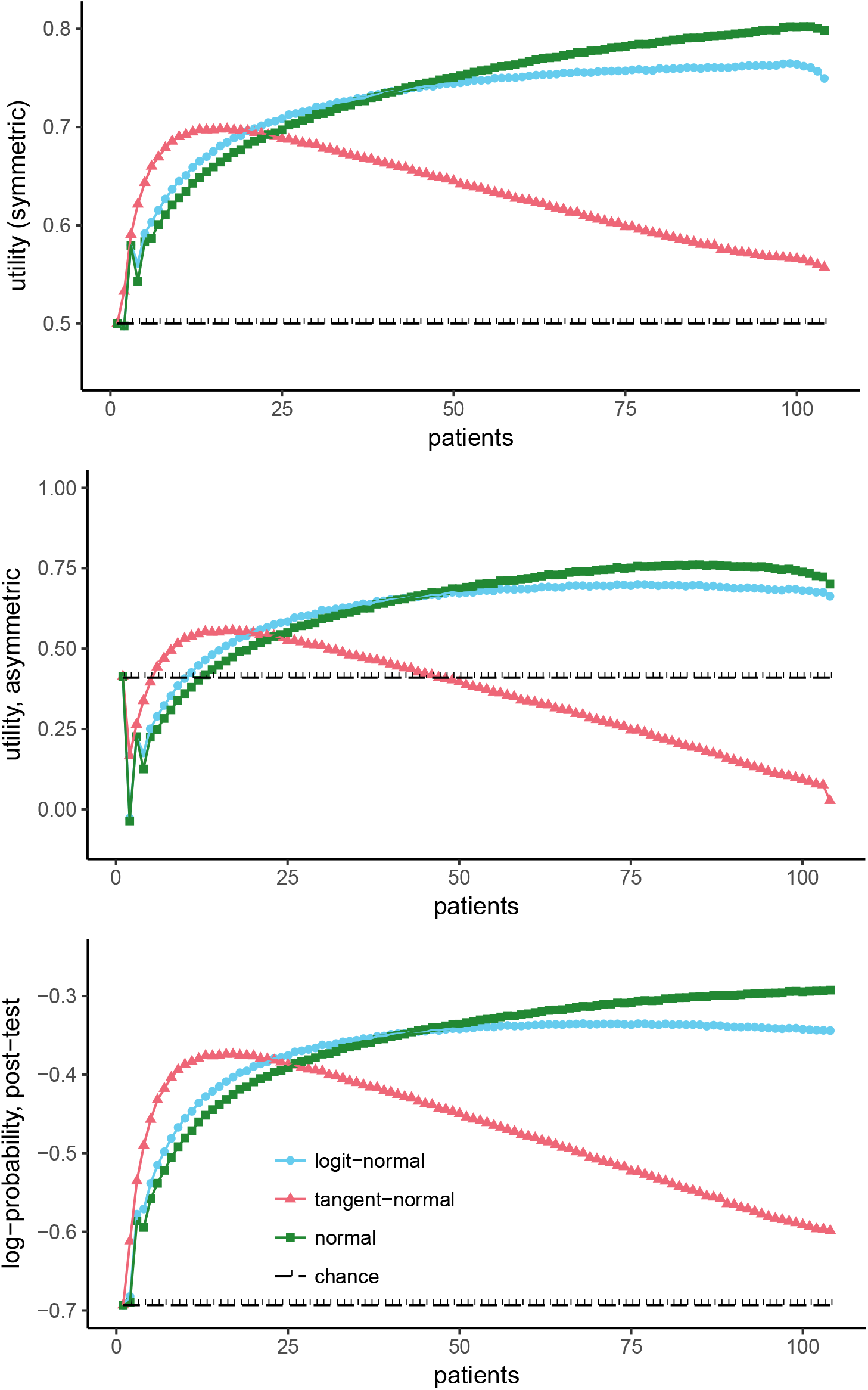
Averaged sequence of utilities, with utility tables (26) and (27), and of log-probabilities for our three models. The standard deviations of the averages are smaller than the markers’ size. The average values for the first and last patients are exact.

**Table 2.** Final results for the three models

|  | symmetric utility (26) | asymmetric utility (27) | log-probability |
| --- | --- | --- | --- |
| normal model | 0.80 | 0.70 | −0.29 |
| logit-normal model | 0.75 | 0.66 | −0.34 |
| tangent-normal model | 0.56 | 0.03 | −0.60 |
| chance | 0.50 | 0.41 | −0.69 |

The plots of fig. 3 illustrate the points made at the beginning of §2.5.1: one model can initially learn faster than another and yet be overtaken in the later stages of learning; this is the case for the logit-normal and normal models. The tangent-normal model shows strong unlearning; this means that a tangent-normal likelihood and its sufficient statistics do not distinguish well between healthy and schizophrenic conditions. The slight downward bends at the final stages of the logit-normal and normal models raise the suspicion that they might also show some unlearning if further training data were supplied.

The trends of the logit-normal and normal models suggest that the learning phase is not finished: more patients are needed before their predictive probabilities become stable. This is also evident from the updated marginal distributions of their parameters (**λ**_H_, ***Λ***_H_; **λ**_S_, ***Λ***_S_), for example those for the connectivity weight *f* between the left superior parietal lobule and left lingual gyrus, shown in fig. 4 for the logit-normal model. The distributions of the location parameters (**λ**_H_, **λ**_S_) have reached the empirical means of the data, but those of the scale parameters (***Λ***_H_, ***Λ***_S_) are still very far away from the empirical variances. The reason is that the prior for the scale parameters had a peak at a very large value of ***Λ***æ 20. The 55 data for healthy patients and 49 for schizophrenic ones have shifted this peak to ***Λ***_H_ = 1.3 and ***Λ***_S_ = 1.5, but more data are needed to shift these peaks to even smaller values – provided that in the meantime the empirical values do not change too much as new data are gathered.

**Figure 4.**
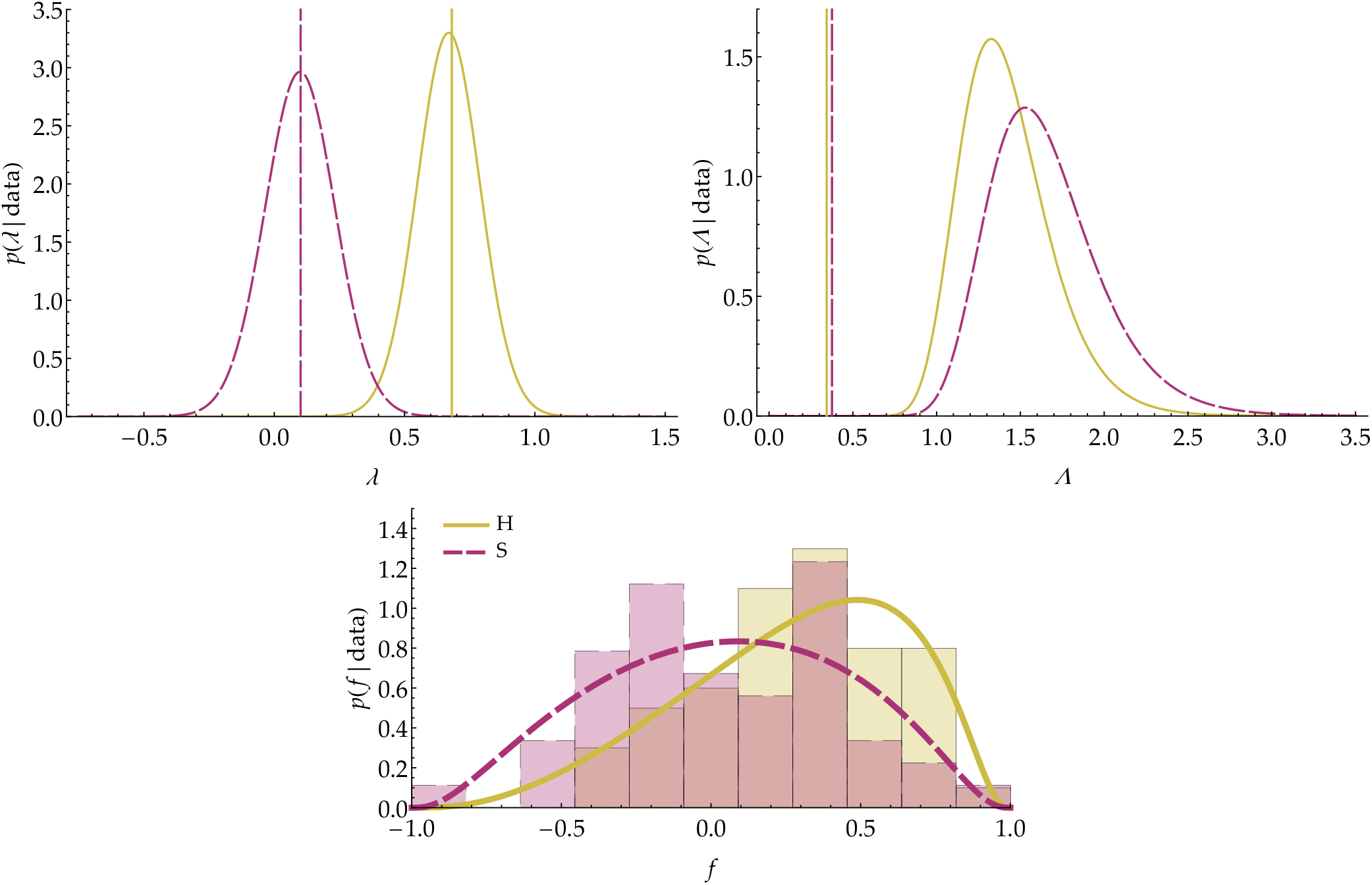
Updated distributions of the logit-normal model for the location parameters (***λ***_H_, **λ**_S_) (left), scale parameters (***Λ***_H_, ***Λ***_S_) (right), and connectivity weights *f*_H_, *f*_S_ (bottom, superposed on the empirical distributions) for the connectivity between left superior parietal lobule and left lingual gyrus, corresponding to the bottom left panel of fig. 1. The vertical lines in the first two plots indicate the corresponding empirical statistics from the data.

The peak at high values of ***Λ*** is a known inconvenient feature of the normal-inverse-Wishart conjugate prior, related to the correlation between correlation and variance components of ***Λ*** characteristic of this prior (e.g.Barnard et al., 2000).

#### 2.5.3 Contrast with other model-comparison criteria

Common Bayesian model-comparison criteria are based on the joint probability that the model gives to training data; especially its logarithm, called “weight of evidence” or “marginal log-likelihood”, or the ratio of such logarithms, called Bayes factors (Jeffreys, 2003, chs V, VI, A; Good, 1950; MacKay, 1992a; Kass and Raftery, 1995). The simple reason is Bayes’ theorem:

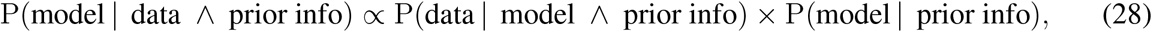

the latter probability usually assumed the same for all models (but see Porta Mana, 2017). A higher weight of evidence means that the model is more probable.

In our study, however, we have two kinds of data: health conditions and fMRI results. Since the likelihood for the health condition, used by the clinician to arrive at a post-test probability, gives the probability for the fMRI results {***f**_i_*}, it seems intuitive to calculate the weights of evidence of our models based on these data. The result is the opposite of what we obtain with the averaged-utility criterion or any of other three mentioned above. We obtain:

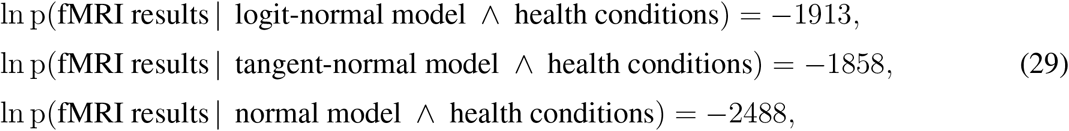

which gives the normal model a much smaller probability than the other two, and the tangent-normal model the highest.

This discrepancy with the averaged-utility criterion is not completely surprising, though. Imagine a disease that leads to no differences at all between the fMRI results of patients with the disease and those of healthy controls. If we found a statistical model that predicted the fMRI results with certainty, this model would thus have a the highest weight of evidence (zero), and yet its final average utility would be at chance level, since it could not help us at all in telling healthy from diseased patients. For our problem the right comparison and selection criterion is the utility or one of the other three criteria previously listed.

#### 2.5.4 Final assessment of models

The average-utility criterion clearly excludes the tangent-normal model, which even shows a rapid unlearning. The logit-normal and normal models have almost similar performances, at around 75–80% of final patients correctly treated. We can also plot the sequence of utilities averaged over healthy and schizophrenic patients separately, as in fig. 5, which gives us an idea of the ratio between true and false negatives, and true and false positives. Both models show around 35% false positives (dismissed schizophrenic patients), with the normal model giving slightly higher final rates of true negatives, 93%, and true positives, 65%, than the normal, 85% and 63%.

**Figure 5.**
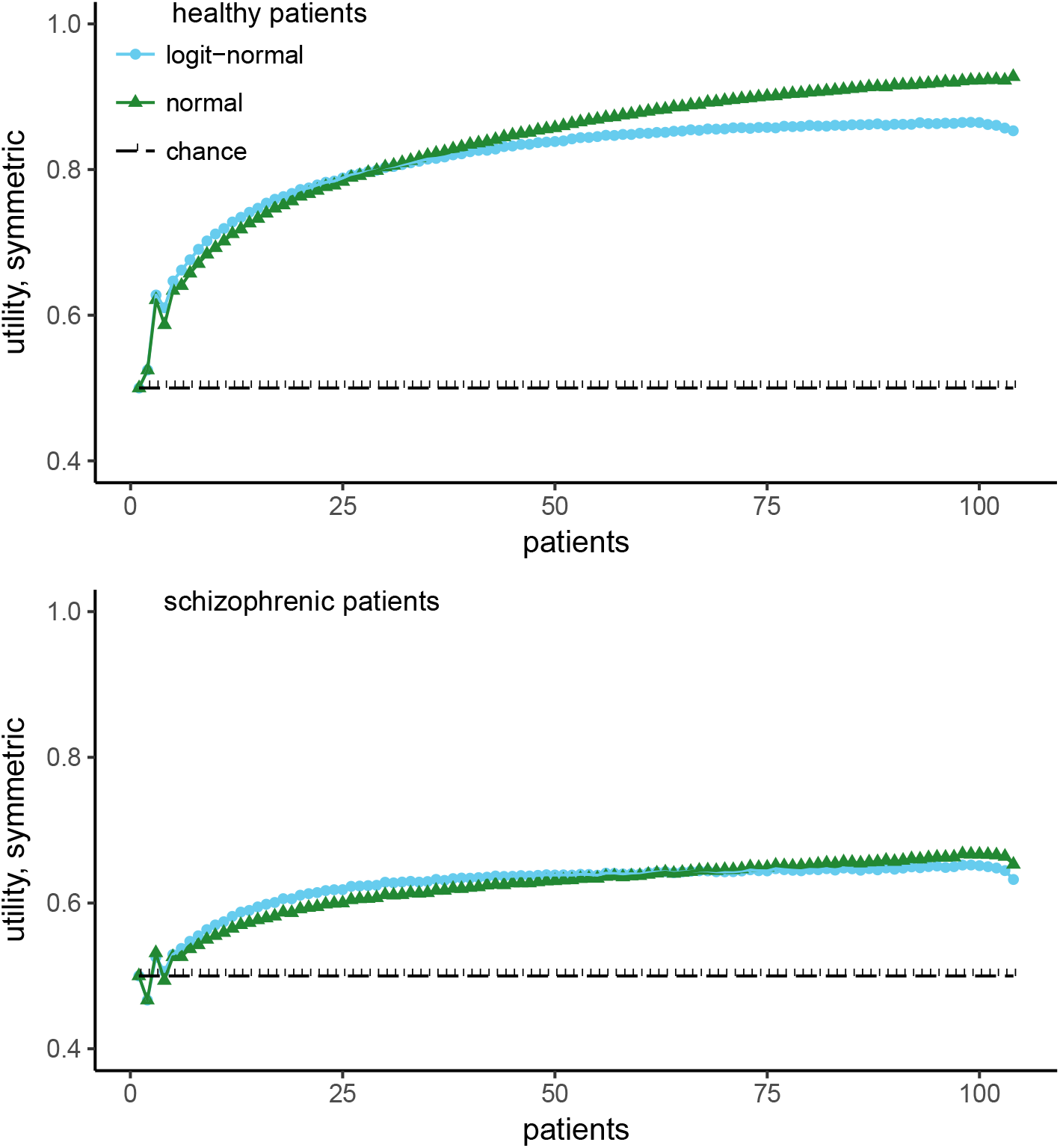
Sequence utilities averaged over healthy (left) and schizophrenic (right) patients separately, for the logit-normal and normal models

However, we emphasize that the features used as inputs to the model were selected using a very simple heuristic (maximum difference in means between the two groups, see §2.3), and the statistical model was selected for its computational properties rather than its fit to the distributions of connection weights derived from fMRI data. This notwithstanding, we believe that with further training, the logit-normal model could reach a higher predictive power. The reason is that some empirical distributions of connectivity weights, like the one for schizophrenic patients shown in red in fig. 4, seem to be bimodal; and the logit-normal likelihood, unlike the normal one, is capable of bimodality, thus fitting these distributions better.

Our assessment, however, is just an illustrative example for the general method discussed in this paper, and we are not earnestly proposing the logit-normal model (nor any other specific model) as the optimal one to use in the problem of diagnosing schizophrenia. We note that any model assessment and selection using the average-utility metric depends on several important quantities and assumptions:

A. the pre-test probabilities given by the clinician; we assumed these to be (0.5, 0.5);
B. the clinician’s range of decisions; we assumed it to be simply “treat or dismiss”, but it could comprise several different kinds of treatments;
C. the utilities of the clinician’s decisions; we assumed these to be as in formula (37);
D. the ratio between the numbers of healthy and schizophrenic training data; 55/49 = 1.12 in our case;
E. the clinician diagnoses one patient at a time.

For a proper model assessment we should therefore investigate and consider more realistic rates of healthy vs schizophrenic cases that visit a particular clinician, in order to have better-informed pre-test probabilities; and we should consider more realistic decisions available to the clinician, together with realistically quantified utilities.

Assumption E. deserves some explanation as it may mistakenly appear that it doesn’t matter whether patients visit the clinician simultaneously or one at a time. Suppose two patients, Tom and Joe, visit the clinician together, and the clinician obtains fMRI data for both, ***f***_T_ and ***f***_J_. The joint post-test probability for their health conditions *c*_T_ and *c*_J_ is different from the one obtained first calculating Joe’s one, say, and then Tom’s using Joe’s results:

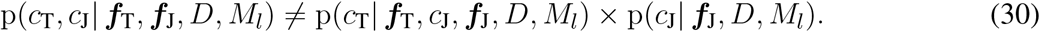

This inequality can be easily verified by applying the probability product rule to the left side, and can be understood as follows. Suppose the clinician first wants to calculate the likelihood for Joe’s being healthy. If Tom is schizophrenic then his fMRI result is unimportant for Joe’s likelihood, owing to our assumption of independent priors (16). But if Tom is healthy, then his fMRI result, *which is known to the clinician*, should lead to an updated model for Joe’s likelihood. The likelihood for Joe’s being healthy is therefore a mixture of these two possible likelihoods, with weights proportional to the post-test probability for Tom’s health condition. Thus, Joe’s likelihood is affected by Tom’s fMRI result even if Joe’s is calculated first and Tom’s health condition is not yet known. More generally, if several patients visit the clinician simultaneously, she should order diagnostic fMRI tests for all of them at once and calculate a joint post-test probability for them, in order to make the best-informed prediction for each.

## 3 DISCUSSION

### 3.1 Summary

The diagnosis of a medical condition is a complex process that takes in a variety of judgements and evidence from the clinician and from any diagnostic tests available to her. Bayesian probability theory has found wider acceptance in medicine because it can consistently combine and frame such judgements and evidence (Goodman, 1999; Davidoff, 1999; Greenland, 1998). Formulae (1)–(3) show the basic scheme of how the clinicians’ judgements and the results of diagnostic tests are combined (Sox et al., 2013). The role of a diagnostic test is not simply to give a dichotomic answer, e.g. healthy/ill, but a *likelihood* for each health condition, to be combined into this scheme together with the likelihoods from other tests. The final probability obtained from these likelihoods is finally used by the clinician to decide upon a course of action, e.g. dismiss/treat (§4.3).

The values of likelihoods from such tests need to be determined from a set of training set of data for each health condition. In this work we have discussed how to determine the likelihoods when the diagnostic test and training set consist of fMRI data, considering for concreteness the case of schizophrenia as the disease in question, and by using real fMRI data from healthy and schizophrenic patients (§2.1). We derived them step by step from first principles through a sequence of assumptions:

a. **Partial exchangeability** with respect to the health conditions, explained in §2.2. We believe this assumption to be very natural in medical diagnostics. By itself it already leads to a specific expression for the likelihood, eq. (8), although this expression is very difficult to compute.
b. **Sufficiency** of an empirical statistics of a reduced set of fMRI data (§§2.3–2.4). We believe such kind of assumption to be sound and at least approximately true when neurologically motivated, and moreover it provides a bridge between biophysical considerations and the specification of probabilities. It leads to a likelihood, eq. (11), amenable to numerical or analytic computation.
c. **Prior knowledge** of the empirical statistics for the different health conditions (§2.4.4). An assumption of this kind is always necessary, especially with small training data sets; but it affects our inferences less and less as our training data accumulate.

In our study, for (b) we specifically assumed the sufficiency of the first and second moments of some functions (§2.4) of the functional-connectivity weights obtained from the fMRI data (§2.3). Our focus on connectivities was neurologically motivated, but our choice of first and second moments of particular functions was made for mathematical simplicity, in order to illustrate the method. Our choice of roughly flat, conjugate priors for assumption (c) was also made for the sake of mathematical simplicity and illustration (§2.4.4).

Notwithstanding our simple, mainly illustrative choices, we obtained good diagnostic performances, briefly discussed in the next section. To assess these performances and the relative predictive power of different choices in assumption (b) above, we presented several criteria, based on decision theory (§2.5). These criteria also help in understanding whether the training of the likelihoods has stabilized (§2.5). We observed that in this kind of study decision-theoretical criteria can yield results in seeming contrast with Bayesian model-comparison criteria, like weights of evidence and Bayes factors (§2.5.3); but this contrast is understandable and is not a sign of inconsistency.

We emphasize that the analysis and conclusions presented here are just illustrations of the general method, and are not meant to be used for real diagnoses: as explained in §§2.4.2 and 2.5.4, for a real application we would first need to investigate better-informed sufficient statistics, realistic decisions available to the clinician, the latter’s utilities, and realistic pre-test probabilities. In this study we used simple default values for illustrative purposes only.

### 3.2 Comparison with other studies and methods

In this study we used a rather naive approach, explained in §2.3, to select a subset of brain connections for our analysis. That approach can be criticized in two different ways. First, it ignores the fact that if distributions are narrow, the means can be close together without much overlap, and that such distributions are likely to be more informative for the purposes of discrimination. This could be improved, for example, by taking the minimal area of the distributions’ overlap (dark red area in fig. 1) instead. Second, we did not restrict our search for suitable connections and associated brain areas to the areas that are known to be part of resting-state networks identified in previous studies, like the default mode network. Our lack of specificity, though, was motivated by previous studies which have demonstrated that functional connectivity can be also measured in other task-related networks, induced by spontaneous activity (He et al., 2009), and that in resting state different activity patterns can appear in schizophrenic patients, owing e.g. to hallucinations during the scan (e.g.Shergill et al., 2000).

Despite our simplified and possibly unrealistic choices of connectivity weights, sufficient statistics, parameter priors, pre-test probabilities, and utility functions, we obtained a diagnostic performance of 80%, measured by leave-one-out cross-validation (§2.5), comparable to classification results based on fMRI data reported in previous studies (e.g.,Venkataraman et al., 2012, 18 healthy + 18 schizophrenic patients; Cetin et al., 2016, 45 + 46 patients; Demirci et al., 2008, 36 + 34 patients).

Our results also compare with classification results that we achieved using machine-learning methods. Cross-validation tests using support-vector machines (using 80% of the data for training and 20% for testing in a randomized iterative way) also yielded around 80% of correct diagnoses (data not shown).

But, as explained in the Introduction (points I., II.), methods like these, which simply classify or are deterministic, do not fit the clinician’s decision-theoretical problem: they cannot be combined with other diagnostic tests and do not fit a general decision-theoretic approach – cf. #x00A7;§2.5.1, 4.3. Most machine-learning methods (Bishop, 2006; Murphy, 2012) are thus ruled out.

There is no real contrast, however, between machine-learning methods and the method presented here: machine-learning algorithms can be interpreted as convenient, fast approximations of Bayesian methods (see e.g. the explicative image in Huszár, 2017), often combined with default utility functions and decision rules (Murphy, 2012; MacKay, 2003, 1992a,e,d,c,b). A machine-learning classification algorithm that gives a good performance can suggest good likelihoods or parameter priors to be used in a statistical model. For example, the simplest version of a support-vector machine (Bishop, 2006, ch. 7; Murphy, 2012, § 14.5) can be interpreted as a model where the likelihood *L*_H_ (***f*** | ***θ***_H_) for one health condition is very large in a half of the dataspace ***f*** ∈ [−1, 1]^d^ and zero in the rest, the hyperplane separating these half-spaces being determined by the parameter ***θ***_H_. The likelihood *L*_S_ (***f*** | ***θ***_S_) for the other health condition is likewise large or zero in two half-spaces separated by a hyperplane determined by ***θ***_S_; see eqs (11) in §2.4.1. In this model the parameter prior p(***θ***_H_, ***θ***_S_ | M, I) correlates the two parameters (***θ***_H_, ***θ***_S_) in such a way that the two hyperplanes coincide and the two likelihoods have support on opposite sides. As the model is trained and the parameter prior is updated, eq. (11b), the hyperplane moves in space in a way to maximize the product of the likelihoods. This corresponds to the search of an optimal separation hyperplane by the support-vector machine.

### 3.3 Possible improvements

Besides using more realistic pre-test probabilities, and utility values, the method presented here could be improved in several other respects, especially with regard to assumptions (b) and (c) summarized in §3.1.

In point (b) we assumed that particular connectivity weights are sufficient to distinguish among schizophrenic and healthy patients. These connectivities were calculated as in §§2.3 and 4.1. More sophisticated choices of Regions Of Interests and functional-connectivity measures (Marrelec and Fransson, 2011; Smith et al., 2011; Wang et al., 2014; Gheiratmand et al., 2017; Demirci et al., 2008) or even integration of functional and structural imaging (Michael et al., 2010) could obviously lead to an increased predictive power. A different choice of sufficient statistics could also improve the performance. With increased computational power it would even be possible to use the full set of connectivities rather than sufficient statistics thereof – so-called “nonparametric” models (cf. Zhang et al., 2014, 2016; Nielsen et al., 2016; Kook et al., 2017).

Regarding assumption (c), in §2.4.3 we mentioned that parametric models may lead to probabilities that lack a closed form and are analytically intractable. The models we chose, with conjugate prior, have the advantage of having closed-form formulae, but they also restrict our choice of sufficient statistics and parameter priors. Higher predictive power could be achieved by using other kinds of sufficient statistics, leading to likelihoods that are not generalized normals, or by using non-conjugate priors, e.g. treating means, correlations, and variances independently, as discussed by Barnard et al. (2000). In this case numerical methods are needed, such as Markov-Chain Monte Carlo sampling and integration (MacKay, 2003, ch. IV; Murphy, 2012, chs 23–24). It must be kept in mind, though, that numerical methods can be computationally vastly more expensive than analytic ones. In preliminary studies that led to our present work we considered a couple of statistical models that require Monte Carlo sampling: a truncated normal and a product of beta distributions among them. The calculation of the relevant integrals for these models has vastly higher time costs than for the closed-form models presented here. For example, calculation of a posterior parameter distribution for the truncated-normal model required 17 h on a computer cluster; whereas the corresponding calculation for the models in the present work takes a fraction of a second on a laptop. To assess and select a model for our clinician to use, such integrals need to be computed over and over, as we explained in §2.5.1. The assessment of statistical models that require Monte Carlo methods can therefore lead to months of computation. Further explorations are needed in this direction.

The comparison with support-vector machines sketched at the end of the previous section shows one important assumption of our statistical models: the independence of the prior parameter distributions for the two health conditions: 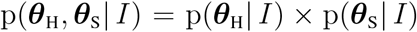, see eq. (16). Statistical models of which support-vector machines are approximations clearly cannot make this assumption. The performance of the model might improve by using a non-independent joint prior distribution, which allows training data for one health condition to influence the parameter distribution for the other. In the case of the generalized-normal models we examined, this can be achieved – whilst preserving their computational convenience – by taking a weighted average of several values of prior coefficients (33). This is known as a hierarchical model (Bernardo and Smith, 2000, § 4.6.5).

The possibilities for improvement listed in these last paragraphs suggest that the method we have presented here, hinging on first principles, has great potential for applications and for development in different directions; moreover this is not to the exclusion of other methods, but assimilating their principles and advantages.

## 4 METHODS

### 4.1 Data preprocessing

Preprocessing of the rfMRI images is carried out using the FMRIB Software Library tools (fsl, v5.08: Jenkinson et al., 2012; Smith et al., 2005) and consists of the following steps: removal of the first ten image volumes, leaving the remaining 130 volumes for further data processing; removing non-brain tissue (Bet: Smith, 2002); motion correction (mcflirt: Jenkinson et al., 2002); spatial smoothing with a 6 mm full width at half maximum normal kernel; temporal low-pass filtering with a cut-off frequency of 0.009 Hz; white matter and cerebrospinal fluid regression (fsl regfilt, melodic).

For each subject we first linearly register the rfMRI image first to the structural, skull-removed image (image segmentation for skull removing with SPM8, Wellcome Department of Cognitive Neurology, London, UKFSL; linear registration with fsl/flirt:Jenkinson and Smith 2001;Jenkinson et al. 2002) and then, through a non-linear mapping, to the MNI standard brain (non-linear registration with Advanced Normalization Tools (ANTS:Avants et al., 2011); MNI 152 standard brain, non-linear 6th generation (Grabner et al., 2006). Regions of interest (ROIs) of the resulting functional image in standard space are extracted such that they match the 94 regions identified by the Oxford lateral cortical atlas with a probability above 50% (Desikan et al., 2006). The temporal mean signals across the voxels in each ROI are used to calculate the functional connectivity measured based on the Pearson correlation coefficient.

### 4.2 The normal model with conjugate prior

This statistical model, denoted in this section by M, is amply discussed in the literature (Gelman et al., 2014, § 3.6; Minka, 2001; Murphy, 2007); here we only give a summary.

Its likelihood is a normal distribution

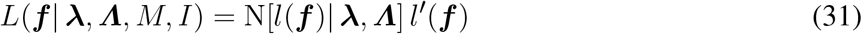

with mean **λ** and covariance matrix ***Λ***.

The prior distribution for the parameters (**λ**, ***Λ***) is a normal-inverse-Wishart distribution, i.e. the product of a normal distribution for **λ** and an inverse-Wishart matrix distribution (Gupta and Nagar, 2000, § 3.4; Tiao and Zellner, 1964; Bernardo and Smith, 2000, § 3.2.5) for ***Λ***:

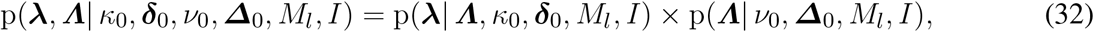

with

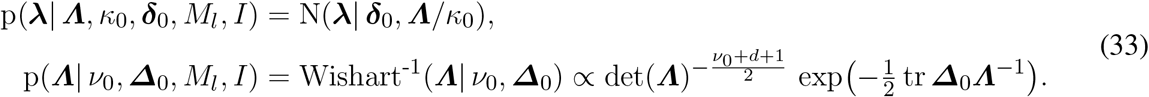

It should be noted how ***Λ***appears as parameter in the distribution for **λ**, so their distributions are not independent. The composite distribution depends on two scalar, one vector, and one matrix coefficients (*κ*_0_, ***δ***_0_, *ν*_0_, ***Δ***_0_).

This prior parameter distribution retains the same form when it is conditioned on the data (***f**_i_*) of *n* patients, becoming a posterior parameter distribution with updated coefficients (*κ*, ***δ***, *ν*, ***Δ***) depending on the prior ones and on the sufficient statistics:

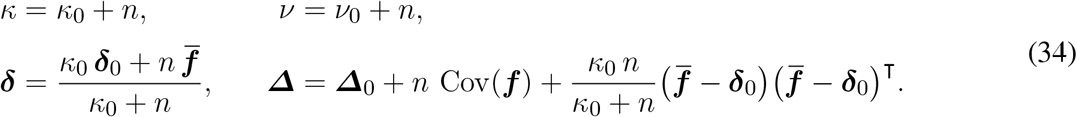

The main features of the normal-inverse-Wishart distribution for (**λ**, ***Λ***) are these:

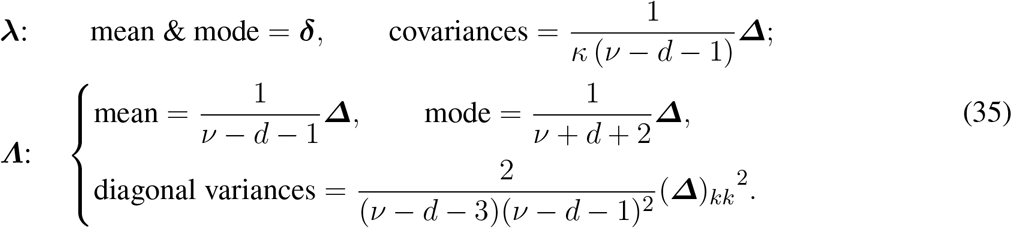

These formulae above say that the uncertainty in the location parameter **λ** decreases as *κ* and *ν* increase for fixed ***Δ***, and the uncertainty in the matrix scale parameter ***Λ*** decreases with increasing *ν*. When *ν* = *d* + 1 the marginal distributions for the correlations of **λ** are uniform (Gelman et al., 2014, § 3.6; Barnard et al.,2000, § 2.2). Because of these properties, a “vaguely informative” parameter distribution should have small *κ*_0_ and *ν*_0_(Minka, 2001; Murphy, 2007).

When the likelihood (31) and the parameter prior (32), updated with (34), are multiplied and the parameters are integrated, the resulting distribution for ***f*** is a multivariate t distribution (Kotz and Nadarajah,2004; Minka, 2001; Murphy, 2007)

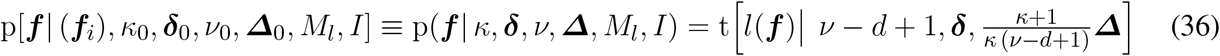

with *ν* − *d* + 1 degrees of freedom, mean ***δ***, scale matrix 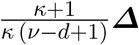, and covariance matrix 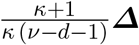.

### 4.3 Decision theory and utility

Once we have the post-test probabilities (*p*_H_, *p*_S_) for the possible health conditions of a patient given the fMRI data, there remains to decide upon a course of action. This is the domain of decision theory (Raiffa and Schlaifer, 2000; Jaynes, 2003, chs 13, 14; Sox et al., 2013, chs 6, 7).

Suppose we have only two courses of action: treat T or dismiss D. A decision-theoretical analysis needs, besides the probabilities for the health conditions, also the utilities (or costs) of choosing an action given the patient’s true health condition. For example, treatment of a healthy patient could harm the latter, or it could be innocuous. With two courses of action and two health conditions we have four utilities *u*_decision|condition_:

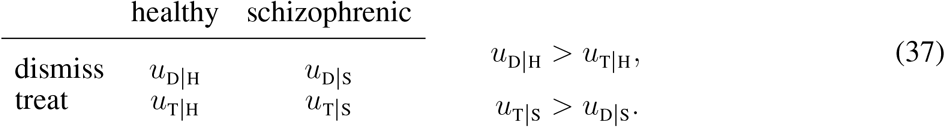

Typically *u*_D|H_ > *u*_T|H_ and *u*_T|S_ > *u*_D|S_, and *u*_D|H_, *u*_T|S_ are positive and *u*_T|H_, *u*_D|S_ negative if we appropriately shift the zero of our measurement units.

The expected utilities for dismissal and treatment are therefore

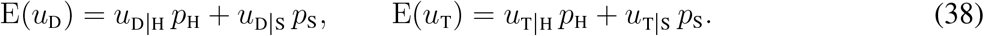

Decision theory says the clinician ought to chose the action having maximum expected utility. For example, she dismisses the patient if E(*u*_D_) > E(*u*_T_), that is if

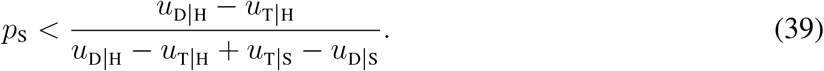

## AUTHOR CONTRIBUTIONS

CB constructed and analysed the graph data from fMRI scans. PGLPM developed the statistical model. They both made the statistical analysis of the data from the model. The manuscript was written PGLPM, CB, AM.

## ACKNOWLEDGMENTS

PGLPM thanks Mari & Miri for continuous encouragement and affection; the kind staff at Iris, where part of this work was done; Buster Keaton and Saitama for filling life with awe and inspiration; and the developers and maintainers of LATEX, Emacs, AUCTEX, Open Science Framework, biorXiv, Hal archives, Python, Inkscape, Sci-Hub for making a free and unfiltered scientific exchange possible. We thank Alper Yegenoglu and Jakob Jordan for support and advice.

## Footnotes

1 http://schizconnect.org/

